# Visuomotor Control of Ankle Joint using Position vs. Force

**DOI:** 10.1101/391367

**Authors:** Amir Bahador Farjadian, Mohsen Nabian, Amber Hartman, Sheng-Che Yen, Bahman Nasseroleslami

## Abstract

Ankle joint plays a critical role in daily activities involving interactions with environment using force and position control. Neuromechanical dysfunctions (e.g. due to stroke or brain injury), therefore, have a major impact on individuals’ quality of life. The effective design of neurorehabilitation protocols for robotic rehabilitation platforms, relies on understanding the control characteristics of the ankle joint in interaction with external environment using force and position. This is particularly of interest since the findings in upper-limb may not be generalizable to the lower-limb. This study aimed to characterize the skilled performance of ankle joint in visuomotor position and force control. A 2-degree of freedom (DOF) robotic footplate was used to measure individuals’ force and position. Healthy individuals (n = 27) used ankle force or position for point-to-point and tracking control tasks in 1-DOF and 2-DOF virtual game environments. Subjects’ performance was quantified as a function of accuracy and completion time. While the performance measures in 1-DOF control tasks were comparable, the subjects’ performance in 2-DOF tasks was significantly better with position control. Subjective questionnaires on the perceived difficulty matched the objective experimental results; suggesting that the poor performance in force control was not due to experimental setup or fatigue but can be attributed to the different levels of challenge needed in neural control. It is inferred that in visuomotor coordination, the neuromuscular specialization of ankle provides better control over position rather than force. These findings can inform the design of neuro-rehabilitation platforms, selection of effective tasks, and therapeutic protocols.

## 1. Introduction

The ankle is a critical joint in human musculoskeletal system contributing to standing, walking, stable balance, and other everyday activities such as driving and pedal operations. Ankle injuries are prevalent problems, in which the neuromuscular control in the joint is affected by the biomechanical injuries such as sprains (Waterman *et al*., 2010), (Waterman *et al*., 2010)) or neural dysfunctions due to stroke or traumatic brain injury (Go *et al*., 2014). The ankle’s role in independent living as well as complex neuromechanical properties of this joint turns the neuro-rehabilitation of the injuries into a high priority and at the same time a challenging task. This has led to numerous motor neurophysiology studies (Goto *et al*., 2014), and in recent years, to the development of novel rehabilitation technologies (Deutsch *et al*., 2001; Farjadian, Nabian, Hartman, *et al*., 2014; Farjadian *et al*., 2015, 2017). While the advances in robotics and automation are providing more efficient and objective rehabilitation experience, they can best be utilized by further understanding of the motor control characteristics of the ankle during interaction with the external environment.

The physical interaction of human foot with the environment is facilitated via the ankle’s force or position control. Ankle’s neuromotor control is a function of neuroanatomical and neurophysiological specialization that are developed by evolution or individual’s lifetime training. Better understanding of ankle joint control properties can enhance the applications of (neuro-)biofeedback (Rockstroh *et al*., 1990; Huang *et al*., 2006; Lünenburger *et al*., 2007), and neuro-rehabilitation (Holden *et al*., 1999, 2005; Sainburg & Duff, 2006). Along this path, we raised the questions: (1) under healthy condition, what are the baseline characteristics of ankle joint force and position control? (2) what are the expected normal performance range of operation? and (3) does human ankle exhibit higher performance over force or position control? Answering these questions would shed more light on the control properties which can serve as a benchmark in neuro-rehabilitation training. Importantly, this information can help to determine what signals (force or position) can be best used in interactive control tasks for (neuro-)biofeedback training (Wolf, 1983; Colborne *et al*., 1993; Bolek, 2003; Femery *et al*., 2004) and neuro-rehabilitation in virtual environments (G. C. Burdea, 1996; Lobo-Prat *et al*., 2014).

We, therefore, sought to answer the question if healthy individuals achieve a better control with force or position, when using their ankle joint for controlling visuomotor tasks, as needed in robotic rehabilitation paradigms or pedal control. This is a non-trivial question, as the hardwired neuro-circuitries in the course of evolution, as well as the plasticity of central and peripheral circuits (due to life experience and practices) both influence the skilled motor performance. From a behavioral perspective, the importance of postural control as a dominant task (involving ankle) may advocate a better position control, whereas the involvement of ankle in dynamic locomotion and running (similar to upper dominant limb) advocates for a better force control.

Several studies have compared the human performance in controlling external targets when individuals used their limb’s force, position or electromyography (EMG). These studies compared the 1-dimensional tracking error, information transfer rate, and other more specific control-engineering measures in upper (dominant) extremity (Corbett *et al*., 2011; Guo *et al*., 2011; Lobo-Prat *et al*., 2014). Overall, force control showed a slightly higher performance over position control in the upper (dominant) limb (Guo *et al*., 2011; Lobo-Prat *et al*., 2014). In these examples, the control superiority was quantified by measures such as lower tracking errors or higher information transfer rates. Surprisingly, however, we could not find a similar study at lower extremity that compares the force and position control in the ankle joint or lower extremity.

Understanding the lower limb characteristics by generalization of findings in upper limb does not seem to be a straightforward task. Some neurophysiological differences between lower and upper limb suggest that the performance of tasks involving visuomotor coordination may be different in ankle: (1) The neuro-circuitries for individuated digit control are evolved differently in lower vs. upper limbs (Hashimoto *et al*., 2013), (2) the H-reflex responses in lower limb are different from those in upper limb (Zehr, 2002), and (3) altering the cutaneous feedback through electrical stimulation affects the performance in visuomotor ankle force-matching, but not the position-matching tasks (Choi *et al*., 2013). Among similarities, the system-level neurophysiological signatures such as motor-related cortical potential are present for both upper limbs (Waldert *et al*., 2009; Nasseroleslami *et al*., 2011a, 2011b; Nasseroleslami, Lakany, *et al*., 2014) and lower limbs (Nascimento *et al*., 2006; Nascimento & Farina, 2008) in position and force control. From a neurophysiological perspective, the complex differences between the upper and lower extremity limit the generalization of the previous findings in upper limb (negligible to some degree advantage for force control) to the lower extremity and ankle joint.

The aim of this study was, therefore, to investigate the characteristic performance of position and force control, in the sensorimotor system at the human ankle joint, using two novel visuomotor game-based tasks. The rationale for this aim was to investigate the skill levels afforded by position and force control that can serve as a prerequisite for the design of rehabilitation paradigms to improve functional outcome of therapy by careful selection of interactive force-or position-based tasks. We designed and conducted experiments to quantify and compare the performance of position and force control in human’s dominant ankle joint at different skill levels. A novel 2 degree-of-freedom robotic system, “virtual reality interfaced ankle and balance trainer” (vi-RABT) (Farjadian, Nabian, Holden, *et al*., 2014; Farjadian, Suri, *et al*., 2014; Farjadian *et al*., 2017), originally designed for actuated assistive/resistive therapy in lower extremity disorders, was utilized for the experiments. To assess the motor control performance during functional tasks, two visuomotor tasks were programmed in virtual environment (a Maze Game and a Board Game). Representative of the range and applications of ankle rehabilitation paradigms (Yoon *et al*., 2006; Farjadian *et al*., 2017), the games were used to provide force or position sensorimotor control challenges to the subjects in 1-D and 2-D inversion-eversion (INEV) or dorsi-plantar flexion (DFPF) conditions. Using the novel robotic system (vi-RABT), performance measures were extracted during human-machine interaction to assess and compare position and force conditions. The objective results collected via the robot were compared against the subject’s perceived difficulty acquired through subjective questionnaires and presented.

## 2. Materials and Methods

### 2.1 Ethics Approval

The experimental design and protocol were approved by the Northeastern University’s Institutional Review Board (IRB# 10-01-12). Subjects provided written informed consent before participation.

### 2.2 Subjects

A total of 27 healthy human subjects (15 females and 12 males, age: 20-40 years) were recruited and participated in the experiment. A screening questionnaire was used to assure the exclusion of subjects with the history of neuromuscular diseases affecting the lower extremity.

### 2.3 Experimental Setup

The experiments were conducted at Northeastern University. The experimental setup for the vi-RABT robotic ankle trainer system (Figure 1) is composed of a robotic footplate, an adjustable chair, a real-time operating machine, the therapist’s station, and the large projection screen. The system is instrumented with force and angular measurement sensors as well as two electrical motors, yielding a “robotic force-plate”. The robot can be locked and used as a static force-plate or alternatively provide controlled actuation to the footplate (Farjadian *et al*., 2017). The free-running operation mode uses active control to cancel the intrinsic inertia of the device in order to provide zero resistance to the subject’s movements about each axis. In this study, we only used the robot in free-running mode (position games) or locked mode (force games) about dorsiflexion/plantarflexion (DFPF) and/or inversion/eversion (INEV) axes of the ankle joint.

**Figure 1.**
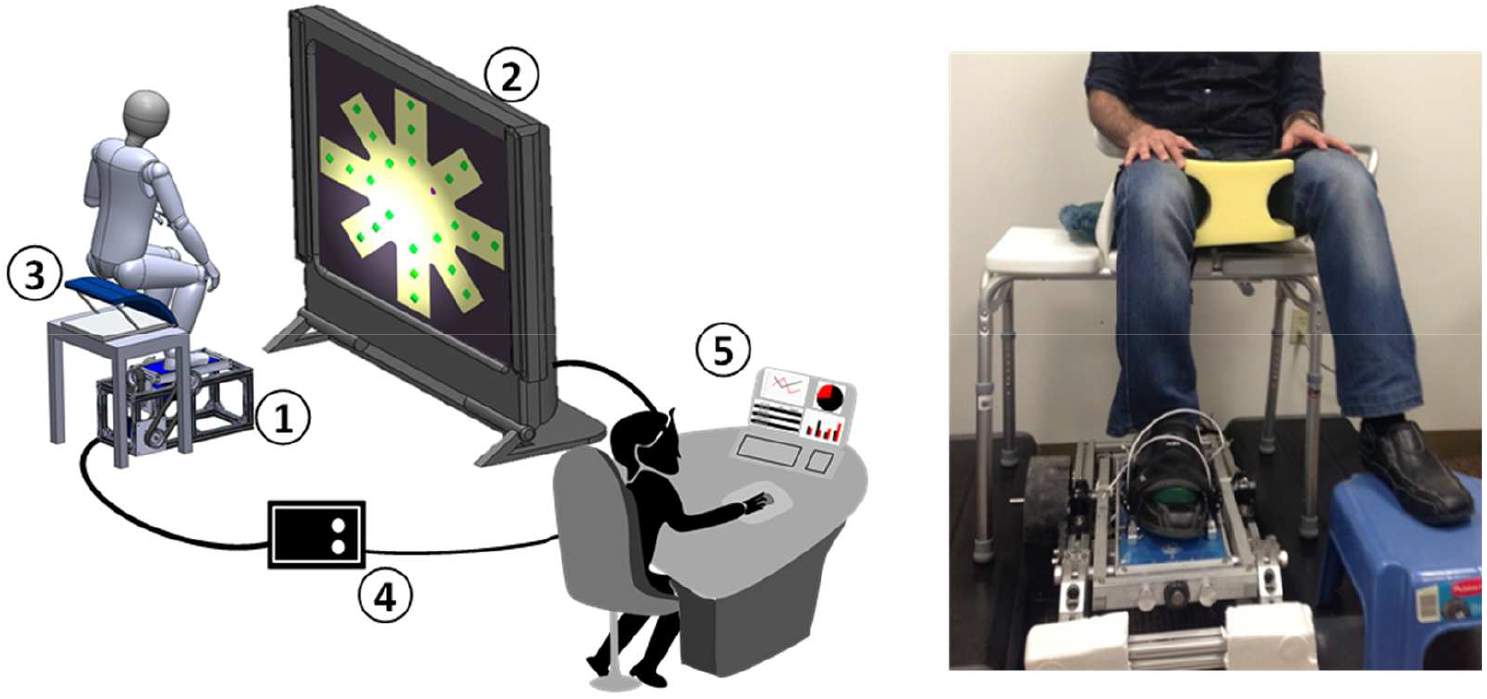
**Left:** The components of the vi-RABT robotic ankle trainer system: (1) 2-DOF robotic footplate, (2) projection screen, (3) adjustable chair, (4) real-time data acquisition and control machine, (5) operator/therapist’s station. The real-time machine controls the subject’s interaction with the footplate and the virtual reality game projected onto the screen. The operator/therapist enters the required parameters and objectively monitors the ongoing experiment and performance. **Right:** Subject is seated on an adjustable chair with the right foot strapped into the robotic footplate. The subject is instructed to control (play) the virtual reality game on the screen via moving the robotic force-plate.

As shown in Fig. 1, the subjects were seated on the adjustable chair, with dominant foot strapped securely into the robotic footplate. To protect subjects’ knee joint and to increase measurement accuracy, subjects’ legs were stabilized with pads and straps to minimize the internal and external rotation of the hip. Additionally, the ankle joint was stabilized in the robotic trainer using heel/calf supports. The chair height was adjusted to place the hip and knee in 90° flexion, and ankle joint in neutral position. The wide projection screen (3×2 m) and the auditory feedback through speakers were used to facilitate subjects’ engagement in the games. The subjects faced the large screen and were encouraged to engage in goal-oriented virtual reality (VR) games and achieve the game objectives. The system acted similar to a joystick for lower extremity that provided the control signals to the VR games, where subjects could play the game on the screen via controlling ankle’s force or position.

### 2.4 Experimental Paradigm

The experimental paradigm was designed to assess the neuromuscular control of the ankle joint in complex and challenging visuomotor tasks, implemented as interactive games in a virtual reality environment. To compare the characteristics of the skilled performance pertaining to neuromuscular system in ankle joint, all subjects performed the games once with force and once with position. In each game, the position of the visual control target was set to be proportional to the ankle’s torque or angular position, for the force or position conditions respectively. Subjects performed the task by their dominant foot (according to self-report) in the beginning of the experiments. These tasks are similar to commonly-practiced daily tasks such as driving and pedal operation, and while not directly comparable to the natural locomotion or posture control, are considered as representative tasks practiced and used in neuro-rehabilitation settings to improve locomotion and posture (Yoon *et al*., 2006; Farjadian *et al*., 2017).

#### 2.4.1 Tasks

Two virtual reality games were used as interactive visuo-motor tasks: A Board Game and a Maze Game (Figure 2).

**Figure 2.**
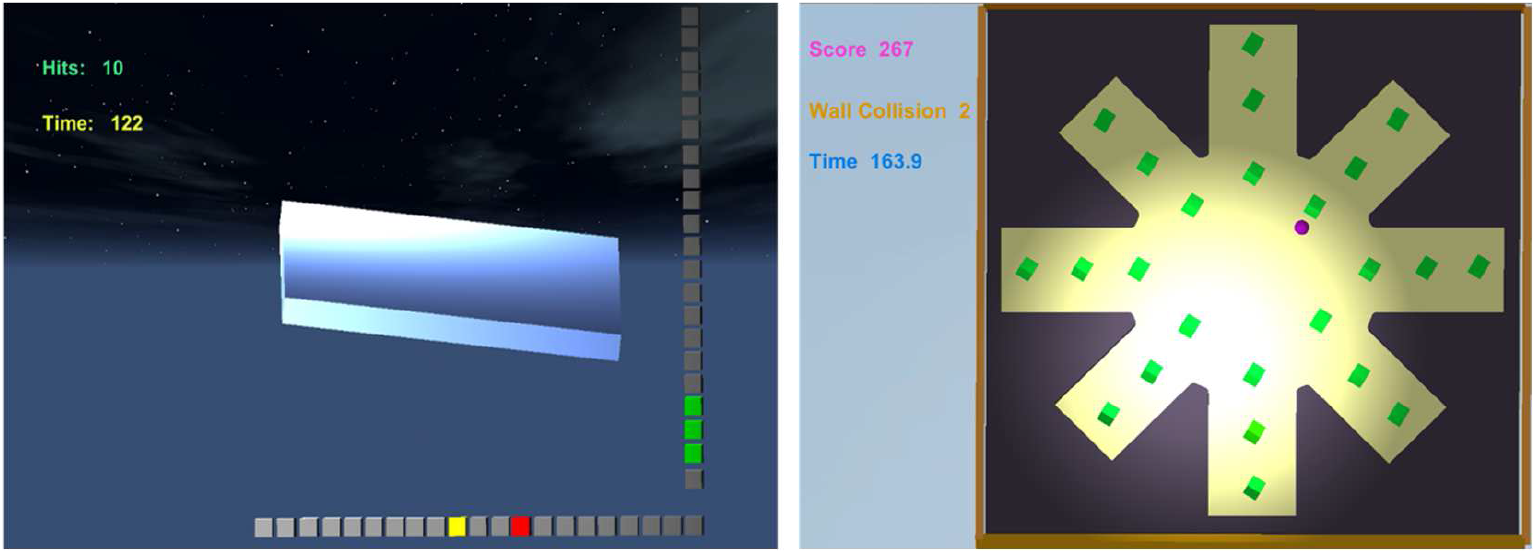
The 2 virtual reality games (subject’s view). **Left: The Board Game.** Subject rotates the virtual board via the robot and achieves the goals. The box lines on the right (Y-axis) and bottom (X-axis) show the current angular position as well as the goal and achievement status about dorsiflexion/plantarflexion (DFPF) and inversion/eversion (INEV) axes, respectively. In this particular example, the subject has achieved the DFPF goal while still looking for the INEV goal on the bottom line. Three highlighted cubes on the right show the achieved status of the DFPF axis. Current position or cursor box (left yellow cube) and next goal (right red cube) are shown on the bottom line. To succeed, the cursor box needs to stay on or next to the target box for 1s. **Right: The Maze Game.** Subject needs to control the moving avatar (purple ball) to acquire and collect all the cubic green goals with minimum collisions to the wall boundaries.

The Board Game consisted of three main components: 1) The blue board (plate), which was used to virtually represent the actual footplate under subject’s foot, 2) The horizontal and vertical box lines, located in the bottom and right sides of the screen, to provide information about angular positions in each axis, and 3) The yellow and red boxes on the box lines to represent the current and desired (target) angular positions, respectively. To succeed in the game the yellow cursor box needed to reach and stay on the red target box for 1 second. Target achievement is fed back to the subject by 3 boxes turning green around the target (see Figure 2 – vertical box line axis). There are a total number of 30 targets that are located following a pseudo-random order. Achieving every target would move the subject to the next target, as shown by new red boxes on the screen. The Board Game was implemented and used in three different scenarios: playing along the y-axis (DFPF) only, along the x-axis (INEV) only, and along both axes (DFPF and INEV) with 2 Degrees of freedom (DOF).

The Maze Game required the subjects to freely move the avatar (purple ball) within a star-shape maze plan to collect all the goals (green boxes). There were a total number of n = 25 goals in the maze surrounded by black walls. The instructions were to collect the goals in the shortest possible time while avoiding wall collision. Collecting the goal was rewarded with a pleasant low-pitch audio signal, while collisions with walls were penalized by an unpleasant high-pitch audio signal.

Subjects controlled the ankle position or exerted torque on the robotic force-plate to play the games. Each subject played a total of 4 games using single or double axes of the ankle joint. This was composed of 3 games in the Board Game: BG-DFPF, BG-INEV, BG-2 DOF and one game with Maze Game: MG-2 DOF. BG-DFPF and BG-INEV were single-axis Board Games along DFPF and INEV axes respectively, and BG-2 DOF, MG-2 DOF were 2-DOF games. The game range (or the maximum cursor’s motion range on the screen) was determined as a function of the subjects’ maximum Range of Motion (ROM) in the movement condition or Maximum Voluntary Contraction (MVC) in the isometric condition. The maximum values for each individual were recorded in the assessment blocks before playing the games.

#### 2.4.2 Protocol

The experimental protocol comprised 3 main components: familiarization, assessment, and game (Figure 3). Starting with familiarization, subjects were given enough practice trials to learn how to interact with the system in different conditions. Both the position and force control paradigms were exposed to the subjects in advance for about 5 minutes, to reduce any potentially confounding factor of learning. Subjects were then assessed in 2 groups (Condition I and Condition II, Figure 3), where the practice blocks either started with force control (n = 20) or position control (n = 7).

**Figure 3.**
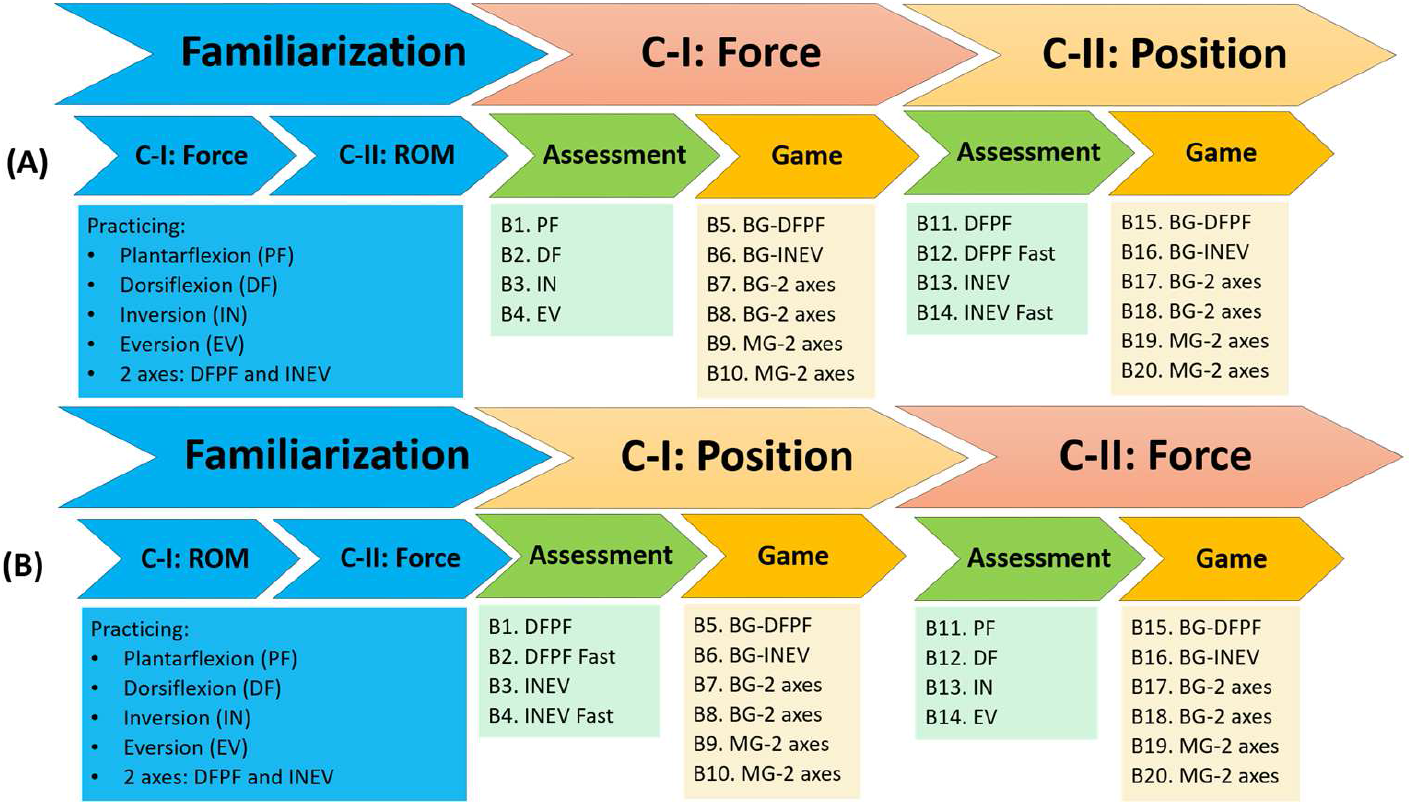
The experimental protocols. After familiarization with the experiments, two groups of subjects (A and B) practices 20 Blocks (B1 - B20) of recording in 2 Conditions (C-I and C-II) with different order of blocks. The assessment blocks (B1-B4 and B11-B14) pertain to the measurement of maximum voluntary contraction (MVC) or range of motion (ROM) in each condition. The game blocks pertain to the interactive board or maze games. BG: Board Game, MG: Maze Game. PF: plantarflexion, DF: dorsiflexion, IN: inversion, EV: eversion.

In the force condition, the assessment blocks were used to measure the MVC when the footplate was locked into the platform and it worked as a force-plate (isometric condition). Subject’s maximum exertion in plantarflexion (PF), dorsiflexion (DF), inversion (IN), and eversion (EV) were measured by having the subjects perform 5 maximal contractions of 3s duration for each of the desired directions. In each case (DF/PF/IN/EV), the corresponding MVC (torque values) was computed by averaging the 5 peak magnitudes. The game boundaries were set as 80% of MVC.

In the position condition, the mechanical locks on the footplate were removed and the controller could make the robot follow the subject’s movements rotating the ankle up, down, in or out. In the assessment blocks, subjects’ maximum ROM was assessed in 7 trials along each of the sagittal and frontal axes at a preferred speed (DFPF, INEV blocks), as well as at the maximum speed (DFPF Fast, INEV Fast blocks). Similar to the force condition, 80% of the maximum ROM values (averaged peak across 7 trials) for each subject was used as the game boundaries. Notice that due to the back-drivable mechanism of the robot, there is negligible external resistance force in this state. Subjects completed playing the same series of games as they did in the force control condition; however, this time controlling the games via angular positions of the ankle.

Taken together, each session included two conditions of force and position measurements. The 10 blocks (Fig. 3) included 190 trials in the force measurements (4×5 trials in 4 assessment blocks, 4×30 trials in 4 board game blocks, and 2×25 goals in 2 maze game blocks) and 198 trials in the position measurements (4×7 trials in 4 assessment blocks, 4×30 trials in 4 board game blocks, and 2×25 goals in 2 maze game blocks). This equals 388 trials in the 20 overall blocks of force and position measurements. A standard rest period of 10 seconds was given after each trial in the assessment blocks, and 15 seconds after each game block. Additional rest periods were given upon individual’s request.

### 2.5 Recordings

The real-time force and position values, as applied by human subjects to the robot, were first recorded by the real-time machine (Figure 1-item 4). In the next step, the data was transferred to the therapist station (Figure 1-item 5) for visualization and storage. The storage sampling rate was 50 Hz.

### 2.6 Evaluation Questionnaires

Upon the completion of each experimental block, subjects completed a Borg’s Rating of Perceived Exertion scale (Borg, 1982) and a Visual Analog Scale (VAS) of the ease of maintaining the balance (of the cursor) during the games (Scott & Huskisson, 1979; Farjadian, 2015). The Borg’s Rating of Perceived Exertion is a measure of the overall task intensity, difficulty and burden perceived by a subject, whereas the VAS is a more specific indication of the cognitive or neural effort required due to the complexity of the game.

### 2.7 Data Analysis and Statistics

The performance of the subjects in the board and in the maze game was quantified by the game completion time (s). In addition, in the Maze Game, the number of collisions (#) reflected the subjects’ accuracy performance. To avoid the potential confounding factor of familiarization and learning effects, we focused on the second block of practice for each game. The paired comparisons for the performance in 1-D conditions and the responses to usability questionnaires were performed using the Wilcoxon’s Signed Rank Test. An analysis of variance (ANOVA) with a within-subject (repeated) design for the force-position condition and a fixed effect for the order of the blocks was used to test the significance of the main effect (force vs. position) and to assess any potential role that the order of conducting the tests may have had. The analysis was performed in MATLAB (version 2017a, MathWorks Inc., Natick, MA, USA). The statistical achieved power was estimated *post hoc* using G*Power (Faul *et al*., 2007).

## 3. Results

All of the subjects successfully completed the experiments. The individuals’ strength, maximum ROM and velocity in the ankle joint were measured in the assessment blocks (Table I).

**Table 1.** The maximum voluntary contraction (MVC) isometric torques, maximum range of motion (ROM) and maximum velocities (V) pertaining to the right ankle joint in each anatomical direction for 27 human subjects (Mean ± SD).

| Direction | Biomechanical Measurements |  |  |
| --- | --- | --- | --- |
|  | MVC Torque (Nm) | Maximum ROM (°) | Maximum Velocity (°/s) |
| Dorsiflexion | 21.36 ± 12.27 | 22.32 ± 5.81 | 69.63 ± 10.20 |
| Plantarflexion | -21.28 ± 11.21 | -26.92 ± 6.01 | |
| Inversion | 7.11 ± 3.45 | 25.78 ± 6.90 | 119.51 ± 21.81 |
| Eversion | -6.07 ± 3.24 | -17.82 ± 6.28 | |

Figure 4 shows examples of the torque and angular position trajectories that a subject used in the board and maze games to interact with and control the virtual environment. Subjects’ torque trajectories showed higher level of fluctuations compared to the trajectories of angular position. Figure 5 (A and B) shows the 2-dimensional representation of the angular positions and isometric torques trajectories, in representative complete trials of the 2-DOF Board and Maze games from a single subject. Figure 5 (B) is the zoomed view of Figure 5 (A) showing the force/position trajectory between two targets. Similar to Figure 4, the trajectories show that in the middle stage of practice, the execution of the game with position control was accompanied by considerably smoother and straighter trajectories, while the execution with force control was accompanied by remarkable fluctuation and non-straight trajectories.

**Figure 4.**
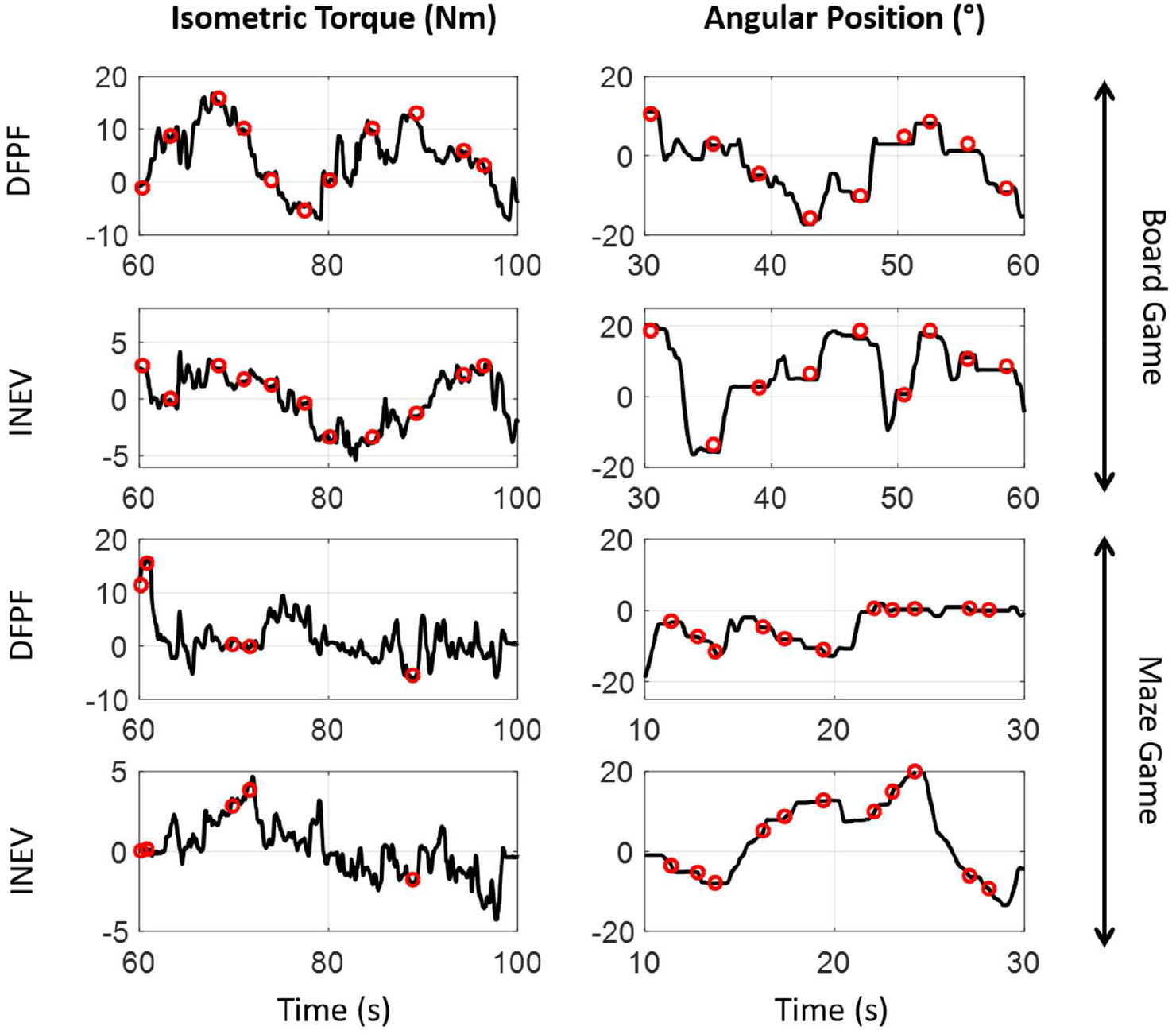
Examples of the ankle joint’s torque and angular position trajectories that a subject used in the board and maze games to interact with and control the virtual environment. The plots pertain to a single trial of the 2-DOF games and the segments in the middle of practice, in 4 anatomical directions. The circle markers represent the target or goal position that the subject matched in the games. Notice the higher level of fluctuation in the control with torque. DFPF: dorsiflexion/plantarflexion, INEV: inversion/eversion.

**Figure 5.**
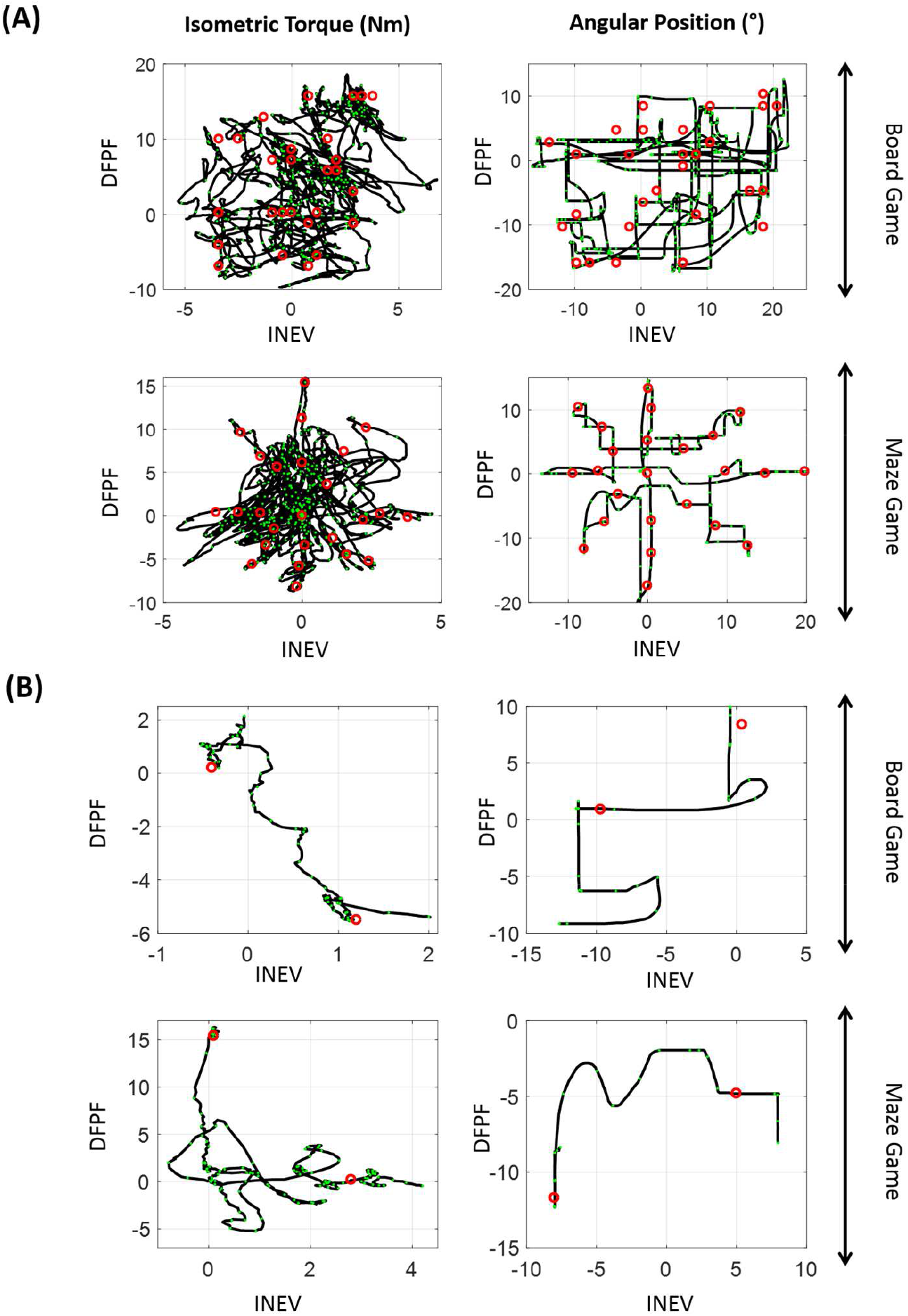
**Top (A):** The 2-dimensional representation of the angular positions (right) and isometric torques (left) trajectories, in representative complete trials of the 2-DOF Board and Maze games from a single subject. The trajectories show that in the middle stage of practice, the execution of the game with position control is accompanied by considerably smoother and straighter trajectories, while the execution with force control is accompanied by remarkable fluctuation and non-straight trajectories. The achieved targets are shown by circles. **Bottom (B):** Zoomed-in view of the torque and position trajectories in (**A**) from a single subject performing 2-DOF Board and Maze games. In each plot, the trajectory between two exemplary targets is shown. The timing information is embedded using the (green) dots (the distances between consecutive dots correspond to 1s) and the achieved targets are highlighted by (red) circles. DFPF: dorsiflexion-plantarflexion, INEV: inversion-eversion.

The trajectories in Figures 4 and 5 imply a more difficult control with force and easier control with position. To more quantitatively compare the performance in the 2 conditions, we compared the game completion times. There was no significant difference between the completion time of the 1-DOF version of the board game (Figure 6) when the same subjects performed them with position versus force control (Wilcoxon’s Signed Rank Test, p > 0.1, n = 27). For the more challenging 2-DOF games, the performance with position control was significantly better for both the board game and maze game (Figure 7). For the Board game, the Force-Position effect showed a significantly better performance (lower completion time) when using position (ANOVA, p = 0.00014, F(1,25) = 20.08, partial η^2^ = 0.45, power = 1-β_0.005_ ~ 0.99) and no significant effect for the order of practice (p = 0.15). For the Maze game, the Force-Position effect showed a significantly better performance when using position (ANOVA, p = 0.0027, F(1,25) = 11.12, partial η^2^ = 0.31, power = 1-β_0.005_ = 0.99) and no significant effect for the order of practice (p = 0.22). A secondary measure of performance in the maze game, the number of collisions (#) (Figure 8), was significantly higher in the force control (ANOVA, p = 0.0054, F(1,25) = 9.45, partial η^2^ = 0.29, power = 1-β_0.005_ = 0.96) with no significant effect for the order of practice (p = 0.50).

**Figure 6.**
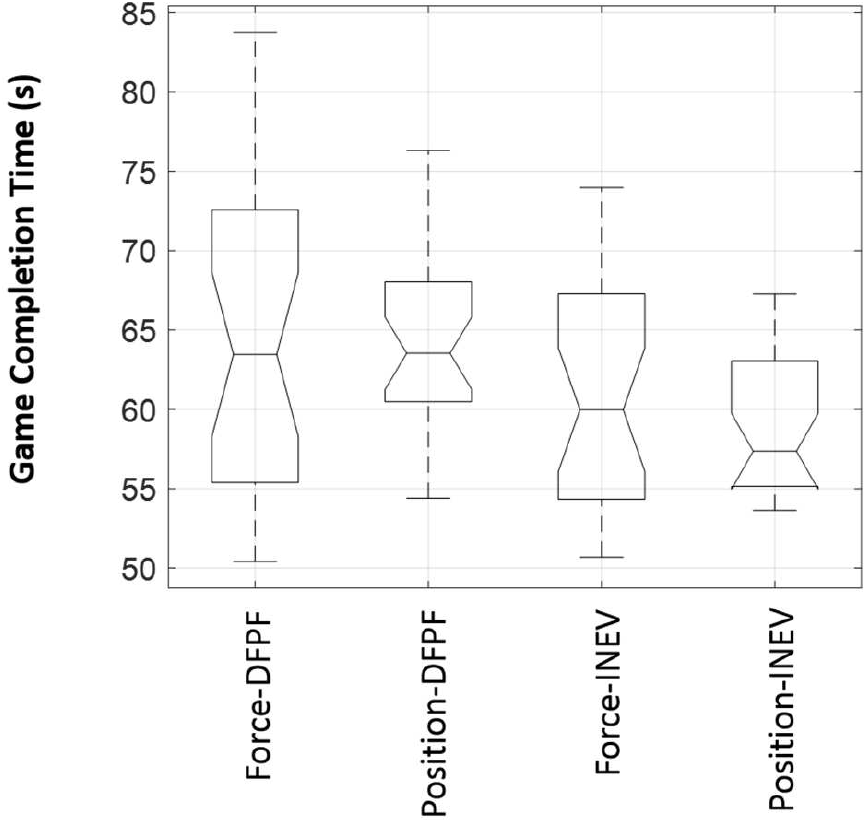
The game completion time in 1-DOF Board Game shows comparable performance with position vs. force control. DFPF: dorsiflexion/plantarflexion, INEV: inversion/eversion.

**Figure 7.**
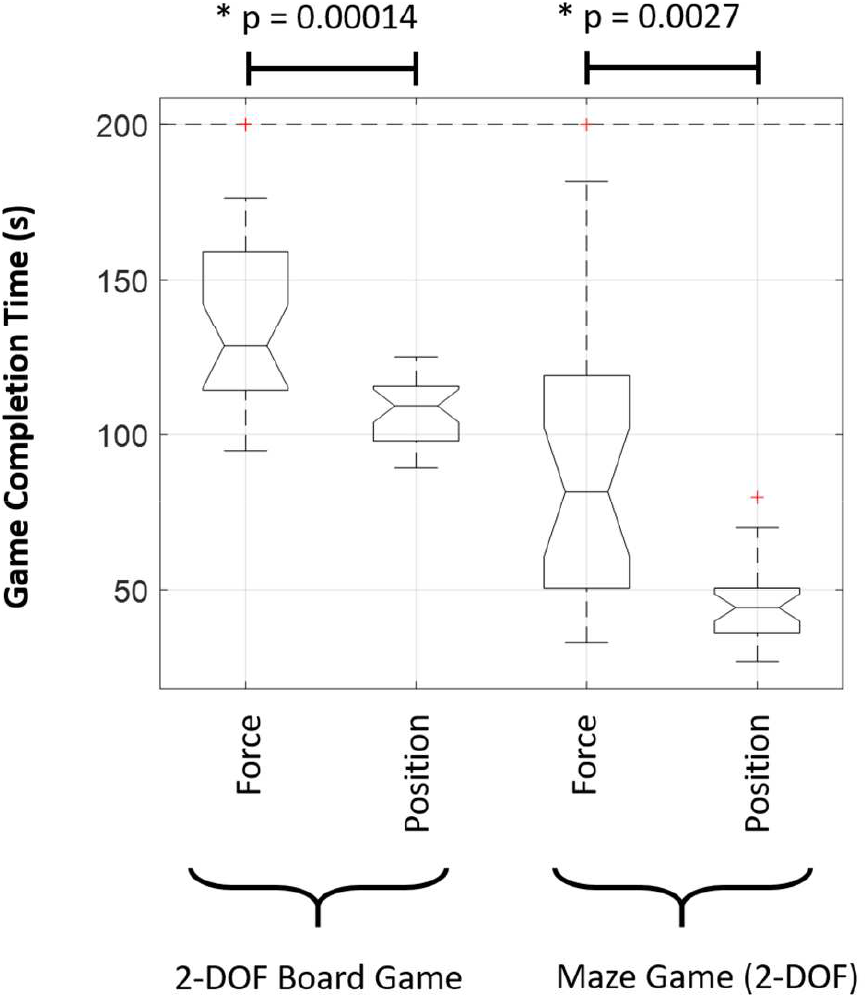
The game completion time in 2-DOF Board and Maze games shows a significantly better control with position control versus force control.

**Figure 8.**
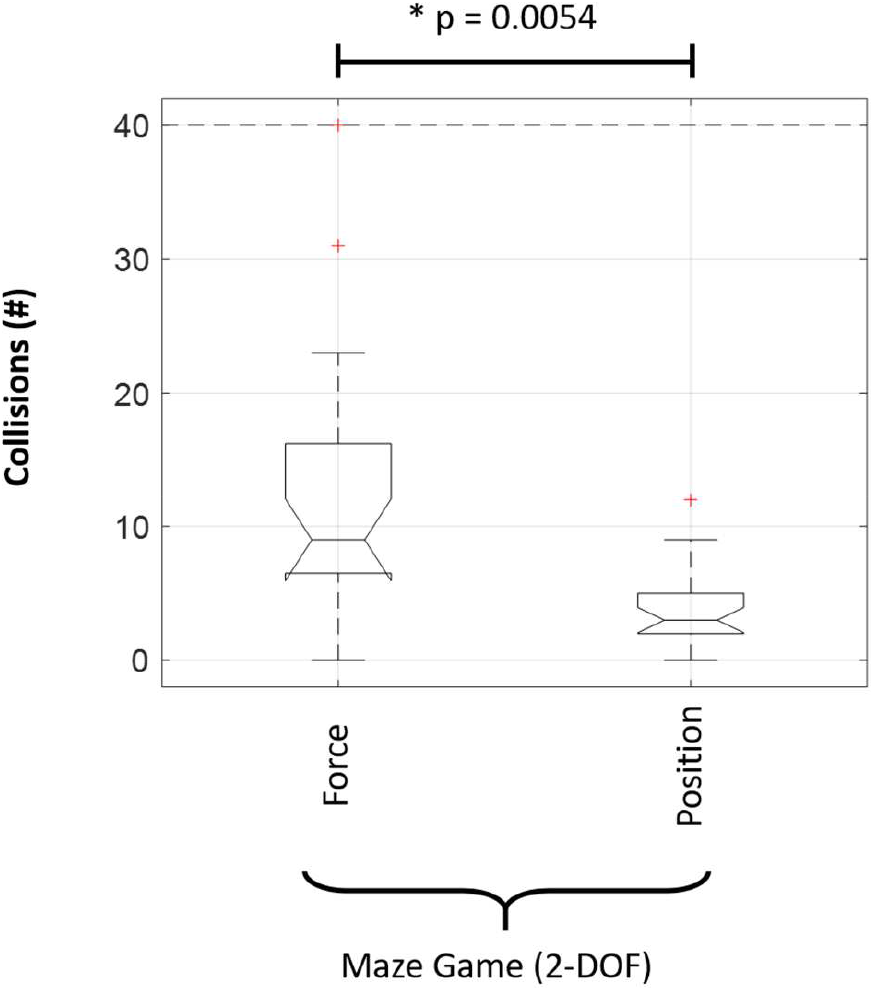
The number of collisions in Maze Game shows a significantly better control with position control versus force control.

Borg’s rating of perceived exertion from 27 subjects (Figure 9) shows that the overall burden experienced by subjects in different games was higher for the force vs. position control (Wilcoxon’s Signed Rank Test, p = 0.000034, p= 0.0002). This holds for the 1-DOF games, as well as the 2-DOF games which had an overall higher level of perceived exertion. The VAS of control (Figure 10) shows that the overall neural/cognitive effort experienced by the subjects in different games was higher for the force vs. position control. This holds for the 1-DOF games, as well as the 2-DOF games which had an overall higher level of neural/cognitive demand (Wilcoxon’s Signed Rank Test for Maze game, p = 0.0037).

**Figure 9.**
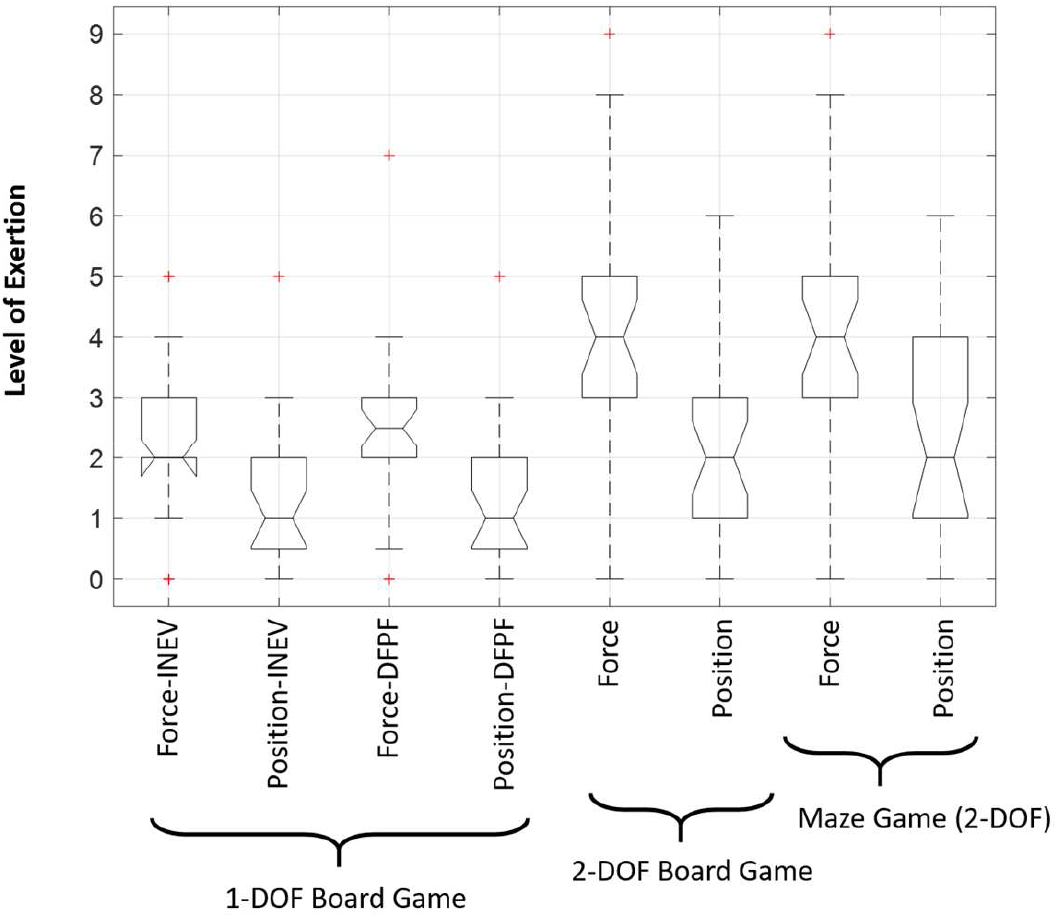
The perceived exertion measured by Borg’s subjective questionnaire was higher for force control vs. position control for both 1-DOF and 2-DOF games. Notice the overall higher exertion intensities for 2-DOF games vs. 1-DOF games.

**Figure 10.**
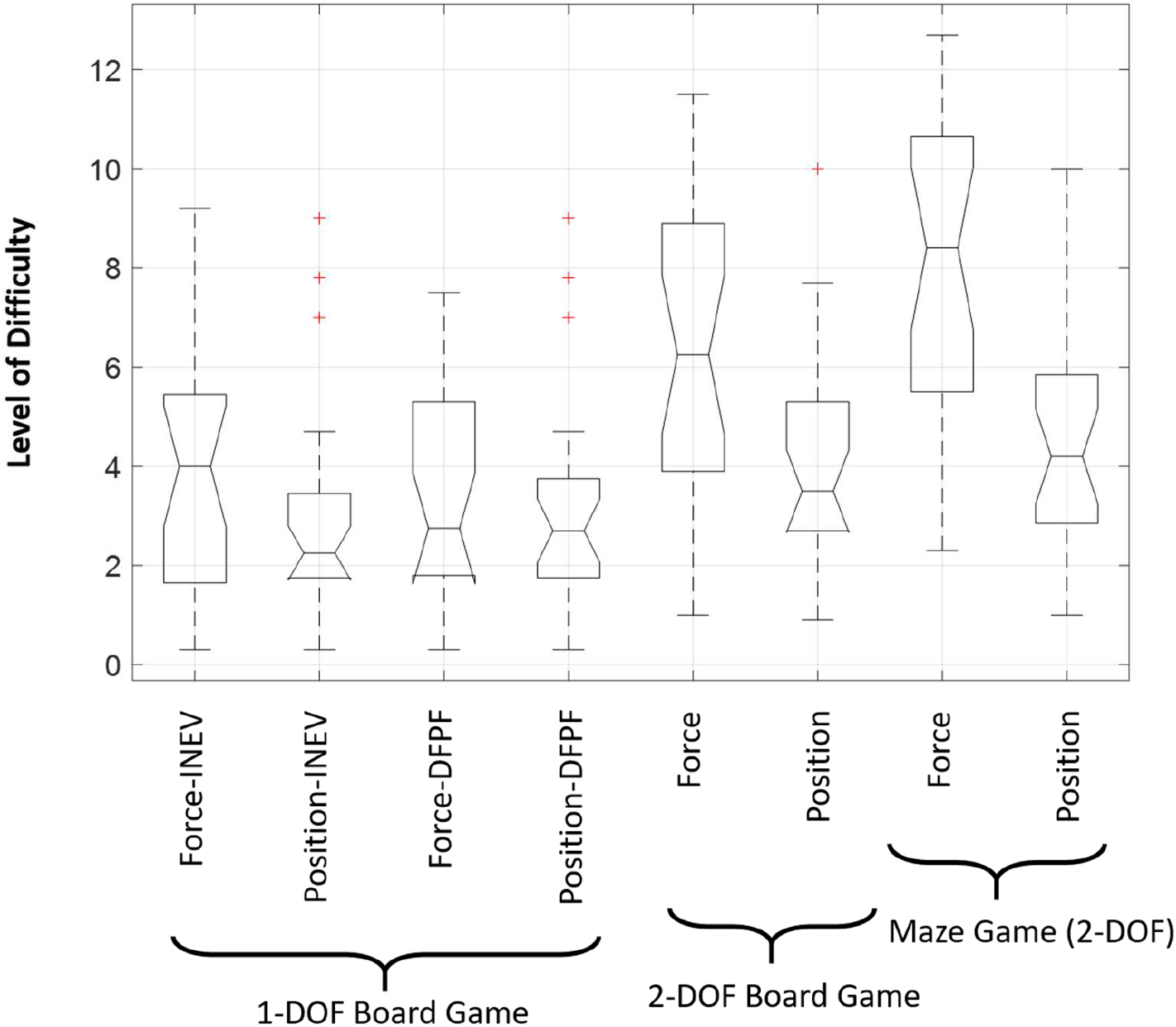
The difficulty in neural control measured by a subjective questionnaire, visual analog scale (VAS) of control, was higher for force control vs. position control for both 1-DOF and 2-DOF games. Notice the overall higher burden of control for 2-DOF games vs. 1-DOF games.

The responses to a usability questionnaire (Farjadian, 2015; Farjadian *et al*., 2017) for the whole experiment that reflected the experienced tiredness (fatigue) showed values only slightly above the neutral level (mean score: 3.48 ± 1.42, 1: disagree, 3:neutral, 5:agree). The expressed cognitive demand showed considerably high values (mean: 4.52 ± 1.05) which was significantly higher than that of mechanical demand (Wilcoxon’s Signed Rank Test, p = 0.00051).

## 4. Discussion

This study was specifically focused on the performance characteristics of skilled visuomotor coordination in ankle joint (rather than upper limb’s) using position and force control. The qualitative comparison of the control trajectories in the force and position control conditions, as well as quantitative inference from the performance measures strongly supports the superiority of performance with position control. This finding differs from the published results on the upper (dominant) limb (Guo *et al*., 2011; Lobo-Prat *et al*., 2014).

The notable difference started developing in 2-DOF games. While the performance in 1-DOF tasks can be considerably affected by the acquired skills for routine daily activities (e.g. pedal works in driving, cycling and sports), the use of 2-DOF games provided a less explored functional tasks for the subjects, and therefore, provided the opportunity to inspect the underlying specialization due to hard-wired neurophysiological circuits and the overall lifetime practice. Moreover, it is important to note that although the completion time (as the performance measure) did not show a significant difference in 1-DOF position vs. force control (Figures 5S–6), the Borg’s (Figure 9) and VAS scales (Figure 10) show that the control was more demanding with force control. Therefore, the more challenging and novel nature of the 2-DOF games served to reveal the underlying differences in the 2 control conditions.

It may be argued that, for a fair comparison, the position and force control experiments need to be mechanically identical in terms of the required level of force. Previous findings on visuomotor object manipulations show that the neural demands can be more critical than mechanical factors in defining the interaction strategies (Nasseroleslami, Hasson, *et al*., 2014; Ye *et al*., 2014). As a result, this study was aimed to emphasize on the task requirements in terms of neural demands rather than mechanical engagement. Nevertheless, the experiments were designed to provide a mechanically equivalent experience pertaining to both position and force control conditions, i.e. each condition covered the 80% of maximum mechanical range (MVC and max ROM), which was a wide range of force and position values according to distributed target positions in the task.

There are other factors that could have biased the findings towards opposite directions (position or force control). Position and force sensorimotor control are reported to show different characteristics at higher range of operation. In isometric torque condition, the exertion levels close to MVC was shown to have higher variability, (Jones *et al*., 2002; Hamilton *et al*., 2004), and similarly the position sense close to maximum ROM may be lower (Darainy *et al*., 2013) than the neutral position. As the position control yielded a better overall performance, the effect of reduced position sense could not be a major factor. We argue against the potential (pure) force demands that could have deteriorated the performance in the force control condition. Comparison of the force trajectories (Figures 4, 5 and 5S) suggests that the fluctuating/noise-looking performance emerge from insufficient ability to control and maintain the force level across the whole range of workspace, and not just closer to MVC. A closer look into the trajectories (Fig. 4–5–5S) pertaining to targets in the center (that need low levels of force for execution) and the borders (that need high levels of force) show comparable patterns of poor control and coordination. Additionally, it is also noteworthy that the performance with 1-DOF force condition (Figure 5S) had a smoother trajectory and seemed less demanding. The 1-DOF experiments can act as a useful baseline or reference showing that the significant differences in the 2-DOF conditions emerged due to the added neural complexity of the task execution rather than mechanical force demand (which did not considerably change from 1-DOF to 2-DOF). Finally, the differences in performance are consistent with the results from subjective questionnaires confirming the difficulty of control with force in terms of the overall perceived intensity and effort (Borg’s scale), but also importantly, in terms of the neural-cognitive burden and demand due to the task complexity (VAS).

The Ia and II sensory pathways from muscle spindles (Pierrot-Deseilligny & Burke, 2012) and the spinal-cord-dominant control, especially for locomotion, as evidenced from animal studies (Prochazka, 1999; Ijspeert, 2008), may be important underlying neuro-circuitries that give rise to the observed specialization for position control. These circuits, though present in the upper limb, may have been overshadowed by the more developed cortical predictive control in the (upper-dominant) limb (Mani *et al*., 2013) leading to a better force control in the upper limb.

It is noteworthy that the “force” and “position” are physical measures specifically of interest in the context of neuro-rehabilitation engineering and as means of quantification for studying human neuromotor system. Indeed, the study of the neuromotor function in terms of these measures does not imply that the sensorimotor system uses these specific measures to encode, decode or control the motor function. The neural signaling in sensorimotor pathways seems to be modulated by a complex function of the combination of force, position and their derivatives (Prochazka, 1999; Mileusnic *et al*., 2006) manifested in neural firing rates. Notwithstanding, the findings can inform of the sensorimotor physiology as quantified by an external observer, which are of practical value for the emerging neuro-rehabilitation technologies.

### 4.1 Limitations

While visuomotor tasks are a common and important group of tasks, further study of other tasks with different sensory and control modalities (e.g. auditory-cued non visuo-motor tasks) will be of interest to test the generalization of the observed performance advantages of position control in the ankle and other joints (e.g. knee and hip). The study included participants in a reasonable age range, nevertheless the elderly people were not included in the study. While a limitation, this could have also excluded the potential effects of aging and the associated age-related neurophysiological and neurological symptoms. The other limitation is the unbalanced sample sizes for the 2 groups of participants (with different order of the tests); however, the statistical analysis took this into account. The analysis did not detect any significant effect due to the order of the experiments and at the same time showed a very high effect sizes (partial η^2^) and the probability of reproducibility (very high statistical power) for the force-position effect. Future studies with additional recordings such as electromyography, EMG (Laine *et al*., 2015; Pizzamiglio *et al*., 2017), electroencephalography, EEG (Nasseroleslami, Lakany, *et al*., 2014; Xu *et al*., 2014; Vuckovic *et al*., 2015), as well as the computational simulation of the potential neural (Nasseroleslami, Vossoughi, *et al*., 2014) and biomechanical (Rashedi *et al*., 2010; Sedighi *et al*., 2011) factors can be used to further elucidate the neurophysiological mechanisms giving rise to the behavioral specialization. Eventually, the findings from this and other studies, need to be assessed with accurate reference to the limb (upper/lower, dominant/non-dominant) and joint (proximal: shoulder, elbow, distal: wrist, hand) involved in the task, and measures used to quantify the skill level.

### 4.2 Conclusion

The subjects’ performance in novel and challenging 2-DOF interactive visuomotor tasks was notably and significantly better with position control, where the poor performance in force control originated from the higher level of neural challenge imposed on the neuromuscular system by force control. The underlying neurophysiological and neuroanatomical specialization of the central and peripheral circuits and the neuromuscular system spanning the ankle joint, as well as the lifetime-learned control can be considered as contributing factors to the advantage of position control over force control. Such characteristic performance differences can be used in design and implementation of neuro-rehabilitation paradigms for more promising therapeutic regimens.

## Acknowledgements

The inspiration and research direction for this study stemmed from the contributions of Professor Constantinos Mavroidis (former advisor to ABF and MN). We dedicate this paper to him, who with great regret is not among us at the conclusion of the study. May his soul rest in peace.

## Author Contributions

Designed the Study: ABF; Performed Experiments: ABF, MN, AH, SCY; Analyzed Data: ABF, MN, SCY, BN; Interpreted Results: ABF, MN, SCY, BN; Prepared Figures: ABF, MN, BN; Drafted Manuscript: ABF, BN; Edited/Revised Manuscript: ABF, MN, SCY, BN; Approved the Final Manuscript: ABF, MN, AH, SCY, BN.

## Data Accessibility Statement

The Data and MATLAB scripts used for statistical analysis are available as supplementary files to the manuscript.

## Abbreviations

ANOVA: : Analysis of Variance
BG: : Board Game
DFPF: : dorsi-plantarflexion (DFPF)
DOF: : Degrees of Freedom
EMG: : Electromyogram
HD-EMG: : High Density EMG
i-EMG: : intramuscular EMG
INEV: : inversion-eversion (INEV)
MG: : Maze Game
MVC: : Maximum Voluntary Contraction
ROM: : Range of Motion
SD: : Standard deviation
vi-RABT: : virtual reality interfaced ankle and balance trainer
VAS: : Visual Analog Scale
VR: : Virtual Reality

## Supplementary Figure

**Figure 5S:**
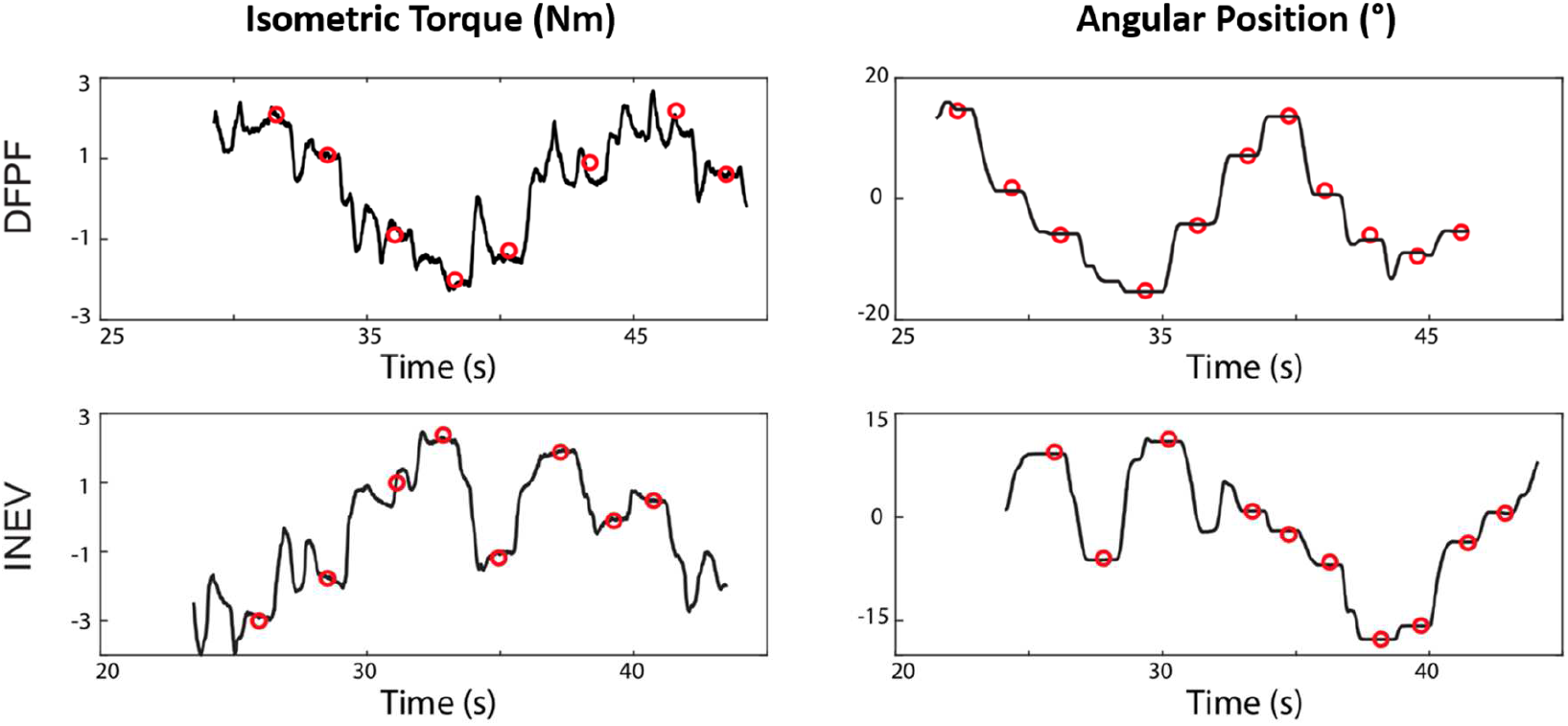
The 1-dimensional representation of the angular positions (right) and isometric torques (left) trajectories, in representative complete trials of the 1-DOF Board game from a single subject. The trajectories show that, in the middle stage of practice, the execution of the game with position control is accompanied by relatively smoother and straighter trajectories. The targets are shown by circles.

